# Copy number variant heterogeneity among cancer types reflects inconsistent concordance with diagnostic classifications

**DOI:** 10.1101/2021.03.01.433348

**Authors:** Paula Carrio-Cordo, Michael Baudis

## Abstract

Due to frequent genome instability and accumulation of mutations during the neoplastic process, malignant tumors present with patterns of somatic genome variants on diverse levels of heterogeneity. The delineation of pathophysiological consequences of these patterns remains one of the main challenges in cancer prognosis and treatment.

Although continuous efforts aim for better characterization of cancer entities through inclusion of molecular characteristics, current ontology systems still heavily rely on clinico-pathological features. Traditionally, malignant diseases have been classified using domain-specific or generalized classification systems, based on histopathological features and clinical gestalt. Aside from the general purpose “International Classification for Diseases in Oncology” (ICD-O; WHO), hierarchical terminologies such as NCIt promote data interoperability and ontology-driven computational analysis.

To evaluate two prominent, general cancer classification systems (NCIt and ICD-O) towards their concordance with genomic mutation patterns we have performed a data-driven meta-analysis of 83’505 curated cancer samples with genome-wide CNA (copy number aberration) profiles from our Progenetix database. The analysis provides a basis to assess the correspondence level of existing classification systems with respect to homogeneous molecular groups, and how individual codes represent an adequately detailed classification.

## Introduction

Malignant neoplasms comprise a group of complex and progressive diseases arising from somatic mutations and with a hallmark of frequently pronounced genomic instability. As individual cancers are composed of populations of cells with distinct phenotypic features and genetic alterations, tumors can be studied through histopathological evaluation as well as mutational analysis. Copy number aberrations (CNA) represent a class of genomic mutations which have been documented in virtually all cancer types and can be assessed genome wide using various methodologies (most prominently through molecular cytogenetic hybridization techniques, WGS [1]. The profiling of CNAs in cancer combined with the study of individual CNA events and genome-wide signatures have been helpful in revealing underlying molecular mechanisms and for the proposal of cancer subtypes [2, 3, 4].

In recent years, and as an addition to or modification of the classifications predominantly based on histopathological characteristics, new cancer entities and subtypes have been established, supported by genomic and transcriptomic analyses of various malignancies. For example, colorectal adenocarcinomas have been separated into CMS1 (microsatellite instability immune), CMS2 (canonical), CMS3 (metabolic) and CMS4 (mesenchymal) subgroups [5]; for medulloblastomas, molecular analysies resulted in a grouping into SHH, WNT, “Group 3” and “Group 4” [6] with only limited overlap with previously defined histological subtypes. The current development in the refinement of cancer type definitions is powered by the addition of molecular parameters as subtype markers or even - as in the case of medulloblastomas - a complete re-evaluation of entity definitions from molecular subtypes with distinct functional mechanisms and clinical trajectories. However, high heterogeneity in cellular phenotypes and dynamic plasticity of tumor microenvironments make tumor categorization a demanding and complicated task with the need to balance between categorical classifications and individual, “personalized” feature definitions. This requires dynamic and coherent classification systems which can allow for iterative expansion and revision of cancer entities and subtypes.

In cancer diagnosis a “classical” pan-cancer classification system is the International Classification of Diseases in Oncology (ICD-O 3) [7] in which a malignant entity is characterized by its morphology (type of cells or tissue that has become neoplastic – kind of tumor – and its biologic activity – behaviour, encoded together) and its topography (site of origin of the neoplasm, encoded as additional term). Over the years ICD-O has been adjusted to recent understandings of tumor biology with revision of some disease areas such as neuroepithelial or hematologic malignancies [8, 9], beyond single code adjustments. However, while the combination of its two coding arms provides very good options for individual disease encodings, the structure of the system has limited flexibility and lacks deeper hierarchical aspects supportive of modern computational concepts.

Increasingly, the use of hierarchical ontologies for biological classifications is being recognized as fundamental for data access, reusability and large scale analysis in the area of cancer research and therapy as well as in other fields of academic biomedicine.

Recent translations and adoptions of hierarchical ontologies with registered identifiers have allowed for internal data structures and novel analysis. As such, important initiatives improving ontology-based search and integration of terminologies are Gene Ontology, MONDO or UBERON [10, 11, 12]. Of special importance in the area of cancer is NCIt Neoplasm Core which provides a controlled vocabulary for specialists at different sub-domains of oncology [13].

In this study, based on the Progenetix collection of human cancer profiles [14, 15], we performed an assessment of popular classification systems (ICD-O 3 and NCIt) through the analysis of inter-sample genomic heterogeneity at different levels of the classification hierarchies based on CNA events.

## Materials and Methods

### Samples and classification systems

A total of 83’505 human oncogenomic profiles (hereforth samples) were retrieved from the Progenetix resource (progenetix.org [16]). The curated samples contained 675 unique NCIt codes, 405 unique ICD-O Morphology codes, 201 unique ICD-O Topography codes, and 1’172 unique ICD-O Morphology-Topography pairs (hereforth ICD-O pairs).

To avoid small sample sizes, the data was then restricted to the 100 ICD-O pairs with the largest number of samples – with all pairs having at least 100 samples each, resulting in 69’769 samples. For each ICD-O pair, samples with CNA parameters indicative of systematic errors or lacking CNV events were removed after statistical clustering of the samples and visual identification of anomalous clusters. This resulted in the removal of 12’491 additional samples, and retaining 57’278.

NCIt provides a morphology-based hierarchy which relates cancers at 9 different levels of detail. The bulk of the unique NCIt codes fall between level 3 and 6 of the tree (as seen in Table 1). There are 134 unique NCIt codes in the sample data, but only 102 of these are contained within the well defined NCIt hierarchy that we study here. 86% of the Progenetix samples correspond to these 102 NCIt codes.

**Table 1.** Number of NCIt codes, and number of Progenetix samples with that code, at the different levels of the NCIt morphology hierarchy.

| Level | 1 | 2 | 3 | 4 | 5 | 6 | 7 | 8 | 9 |
| --- | --- | --- | --- | --- | --- | --- | --- | --- | --- |
| N. of NCIt codes | 1 | 14 | 160 | 331 | 407 | 272 | 116 | 30 | 8 |
| N. of samples | 0 | 0 | 4’662 | 7’156 | 21’677 | 20’972 | 4’578 | 402 | 0 |

We defined a level 2 cutset of a given level 2 NCIT code to be all NCIT codes that are any progeny of it (children, children of children, etc.). There are 14 level 2 NCIT codes, and hence 14 level 2 NCIT cutsets. The filtered ICD samples are centered in specific branches of the NCIT tree: 84 percent of samples belong to the top 4 cutsets, with four cutsets not containing any samples (see Supplementary Material - Table **??**).

### Data representation

As CNA events are common in cancer samples, but with differences in their coverage of the total genome depending on the cancer type -however with great variation and overlap in the statistics-(see Supplementary Material: Table **??**), the study of CNA events needs to look at patterns in how the CNA are distributed across the genome.

The raw data provides individual segments where CNA have been detected. Each sample can have multiple of such segments (with different length and intensity) along the genome, positive for duplication, and negative for deletions. To enable robust statistical analysis, a single vector representation of each sample is constructed through aggregation of the raw CNA events across the chromosomes.

Each chromosome, j = 1,…, 22, is split into *b_j_* bins, with approximately equal length of the bins across the chromosomes, with the desired number of bins pre-specified. Technically, 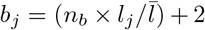 where *n_b_* is the desired average number of bins per chromosome, *l_j_* is the length of the chromosome, and 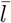 the average chromosome length. Here *n_b_* = 5 was chosen, resulting in a 246 dimensional observation *x_i_*.

Chromosome *j* has 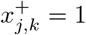 if bin k contains a duplication CNA, and 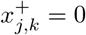 otherwise. Similarly 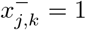 if the k^th^ bin contains a deletion CNA, and 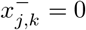 otherwise. For chromosome j for duplications 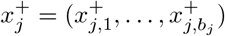, and similarly for deletions, 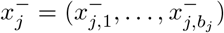. A single sample then has vector 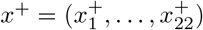 for duplications across all chromosomes, as well as 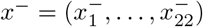 for deletions. Finally, a single sample 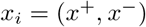 is the concatenation of the duplication and deletion vectors, where subscript i has been added to refer to a specific sample. The length of *x_i_* is therefore 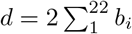.

For CNA events we selected an exclusion threshold (*log_2_* of inferred tumor / reference signal) of 0.1 (or −0.1, for losses), based on empiric values with results showing limited sensitivity to reasonable variations of this value. Next, for called CNA events, the percentage of the bin that is covered by the CNA was computed and bins with a coverage of less than 5% were assigned a negative (0) status.

The binary representation and the selection of a smaller number of bins was found not to degrade the ability to distinguish between pairs of different cancers in supervised as well as unsupervised classification exercises. Further, results were not highly sensitive to the value of the threshold applied.

### Multivariate Bernoulli distribution for binary vectors

To allow for probabilistic modelling of the binary CNA vectors, they were treated as being realizations from a multivariate Bernoulli distribution (MBD). This distribution generates vector random variables *X_i_* = (X_i,1_,…, X_i,d_) with each element *X_i,j_* independently taking the value 1 with probability *p_j_* and the value 0 with probability 1 – *p_j_*. As such, each element can have different probabilities of taking the value 1 (i.e., encoding presence of a CNA in that bin). Each bin was treated as independent. The probabilities were collected in vector *p = (p_1_,…, p_d_)*, which is the CNA profile referred to in the text.

In the event of mixed populations – e.g., some specified fraction of samples come from a MBD with probabilities *p^a^* with the rest coming from an MBD with *p^b^* – then one has a mixture of MBD with two clusters. The probability parameters, as well as the parameters that specify relative weight of each MBD, can then be estimated by an Expectation Maximization algorithm [17]. The model also allows to assign observations to the most likely cluster.

### Clustering & dissimilarity

To assess the quality of a clustering one needs a measure of heterogeneity, which is called dissimilarity. For a set of vector observations *x_i_*, i = 1,…*n* where *x_i_ = (x_i,1_,…, x_i,d_*), the dissimilarity, 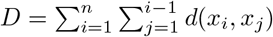, is the sum of all pairwise distances, where d is a distance metric. When the n observations are split into distinct partitions/clusters, the total dissimilarity (computed separately for each partition and then summed) will be less than the case of using a smaller number of partitions – in particular where genuine clusters exist.

A common measure of dissimilarity is *squared Euclidian distance*, 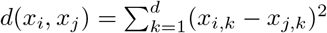. This has the computationally convenient property that the total dissimilarity can be computed as 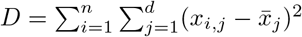 where 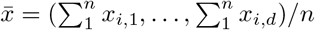 is the *mean centroid*. Note that D is the sum of the *total sum of squares* (TSS) for the d separate dimensions. TSS is simply the sample variance without dividing by the sample size.

For binary valued variables, as used here, applies that the dissimilarity level of a cluster is not immediately meaningful, nor can dissimilarity of different clusters be compared. This is because, as established above, dissimilarity is related to variance, which for binary (Bernoulli) variables is *V_ar_*(*X*) = *p*(1 – *p*) where *p* = *P_r_*(*X* = 1), and as such the dissimilarity depends on the value p. For example, for vectors with *d* = 2, a mixed cluster with two equally weighted MBD: one having *p* = (0.1, 0.1) and the other *p* = (0.9, 0.9) will have the same overall dissimilarity as samples from a cluster coming from a MBD with *p* = (0.5,0.5) – where by definition the latter is less heterogeneous. As such, reduction in dissimilarity by introduction of additional sub-clusters must be used. In the case of the mixed cluster this could be achieved. Also, in this case the squared Euclidian distance is quite general as it is proportional to other common dissimilarity measures such as the l1 norm, and simple matching coefficient.

Note that from the well known decomposition of total sum of squares, the reduction in dissimilarity from the single cluster case (all observations together) to the case of c clusters is also equal to 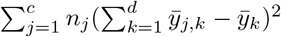 where c is the number of clusters, *n_j_* is the number of observations in cluster j, d is the dimension of each observation, 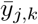 are elements of the cluster-j specific medoid (mean values), and 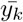 are elements of the medoid for the single cluster case (all observations together). As such, the reduction in dissimilarity is related to the dissimilarity between the medoids themselves. Therefore the greater the reduction in dissimilarity, the greater the initial heterogeneity.

The hierarchical clustering that is performed in this work uses the R *hclust* function, with the Ward scheme [18].

## Results

### NCIt hierarchy assessment

First it was tested if the current level of detail of the NCIt tree was justified. Further testing of the parent-child structure of NCIt and the clustering of samples entailed at different levels was performed. Finally, it was analyzed what further level of detail (sub-clusters) was statistically justified.

### Level of detail in NCIt code mappings

To assess the resolution of the NCIt system, it was tested if the NCIt system separates samples in a way such that the samples form clusters that were statistically different. Specifically, for a given NCIt code, the samples corresponding to that NCIt code or any of its progeny (cutset) in the NCIt tree were used to compute a profile (average CNA probabilities). The profile was then compared with the profiles for all other NCIt codes at that level. This testing was done at all 7 levels of the NCIt tree, where at each level all NCIt codes were pair-wise tested against one another. To ensure sufficient sample size, only NCIt codes with at least 100 samples were retained.

As summarized in Table 2, at all levels multiple pair-wise tests were performed. The test between two clusters was based on the squared distance between the average profile for each cluster, where the p-value was computed by random assignment of samples to clusters. In all cases, all tests found the pairs to be significantly different (p<0.01). On this basis, the level of detail in the hierarchy was justified in all cases.

**Table 2.**
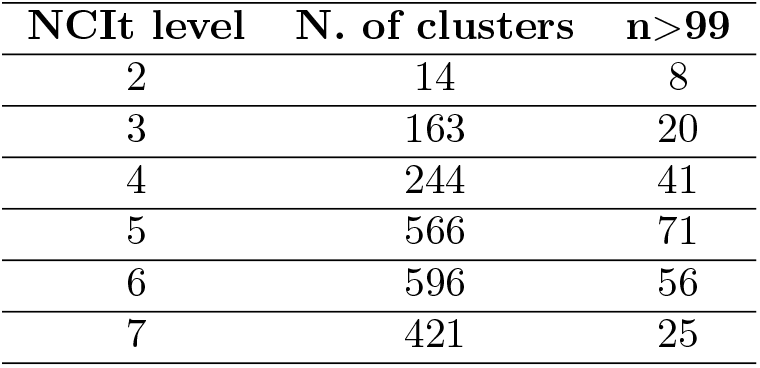
For each NCIt level, the number of clusters, and the number of NCIt codes at that level having at least 100 Progenetix samples.

| NCIt level | N. of clusters | n>99 |
| --- | --- | --- |
| 2 | 14 | 8 |
| 3 | 163 | 20 |
| 4 | 244 | 41 |
| 5 | 566 | 71 |
| 6 | 596 | 56 |
| 7 | 421 | 25 |

**Table 3.** Reduction in dissimilarity for the 30 ICD-O cancer types with the most Progenetix samples. Provided are dissimilarity for the 1 cluster per cancer case (D1), the 4 clusters per cancer case (D4), and the percentage difference.

| ICD-O Code | Morphology - Topography Term | $D_1$ | $D_4$ | $(D_4 - D_1)/D_1$ |
| --- | --- | --- | --- | --- |
| 8140/3 C34.9 | Adenocarcinoma, NOS - Lung, NOS | 116482 | 102843 | 0.12 |
| 8140/3 C18.9 | Adenocarcinoma, NOS - Colon, NOS | 85867 | 75508 | 0.12 |
| 8140/3 C15.9 | Adenocarcinoma, NOS - Esophagus, NOS | 20369 | 17852 | 0.12 |
| 8070/3 C10.9 | Squamous cell carcinoma, NOS - Oropharynx, NOS | 16711 | 14661 | 0.12 |
| 8046/3 C34.9 | Non-small cell carcinoma - Lung, NOS | 64521 | 56427 | 0.13 |
| 8041/3 C34.9 | Small cell carcinoma, NOS - Lung, NOS | 20299 | 17712 | 0.13 |
| 8720/3 C44.9 | Malignant melanoma, NOS - Skin, NOS | 61745 | 53100 | 0.14 |
| 9680/3 C77.9 | Malignant lymphoma, large B-cell, diffuse, NOS - Lymph node, NOS | 27259 | 23313 | 0.14 |
| 8170/3 C22.0 | Hepatocellular carcinoma, NOS - Liver | 29969 | 25651 | 0.14 |
| 9440/3 C71.9 | Glioblastoma - Brain, NOS | 73714 | 62791 | 0.15 |
| 8441/3 C56.9 | Serous cystadenocarcinoma, NOS - Ovary | 63187 | 53878 | 0.15 |
| 8140/3 C25.9 | Adenocarcinoma, NOS - Pancreas, NOS | 25072 | 21340 | 0.15 |
| 8500/3 C50.9 | Infiltrating duct carcinoma, NOS - Breast, NOS | 379631 | 317528 | 0.16 |
| 8140/3 C61.9 | Adenocarcinoma, NOS - Prostate gland | 70322 | 58541 | 0.17 |
| 8520/3 C50.9 | Lobular carcinoma, NOS - Breast, NOS | 21300 | 17774 | 0.17 |
| 9690/3 C77.9 | Follicular lymphoma, NOS - Lymph nodes, NOS | 8397 | 6997 | 0.17 |
| 8070/3 C15.9 | Squamous cell carcinoma, NOS - Esophagus, NOS | 18882 | 15604 | 0.17 |
| 8070/3 C53.9 | Squamous cell carcinoma, NOS - Cervix uteri | 14030 | 11699 | 0.17 |
| 8310/3 C64.9 | Clear cell adenocarcinoma, NOS - Kidney, NOS | 24983 | 20591 | 0.18 |
| 8070/3 C34.9 | Squamous cell carcinoma, NOS - Lung, NOS | 50776 | 41287 | 0.19 |
| 9470/3 C71.6 | Medulloblastoma, NOS - Cerebellum, NOS | 51091 | 40653 | 0.20 |
| 8010/3 C56.9 | Carcinoma, NOS - Ovary | 36042 | 28672 | 0.20 |
| 9500/3 C47.9 | Neuroblastoma, NOS - Peripheral nervs incl. autonomous | 19766 | 15902 | 0.20 |
| 8010/3 C50.9 | Carcinoma, NOS - Breast, NOS | 38605 | 30133 | 0.22 |
| 8140/3 C16.9 | Adenocarcinoma, NOS - Stomach, NOS | 37093 | 29030 | 0.22 |
| 9861/3 C42.1 | Acute myeloid leukemia, NOS - Bone marrow | 23022 | 17700 | 0.23 |
| 8380/3 C54.1 | Endometrioid adenocarcinoma, NOS - Endometrium | 15060 | 11346 | 0.25 |
| 9731/3 C42.1 | Plasmacytoma, NOS - Bone marrow | 32646 | 24166 | 0.26 |
| 9823/3 C42.1 | B-cell chronic lymph. leukemia/lymphoma - Bone marrow | 11307 | 8313 | 0.26 |
| 9960/3 C42.1 | Myeloproliferative neoplasm, NOS - Bone marrow | 5551 | 2515 | 0.55 |

### Assessment of parent-child structure of NCIt

The quality of the NCIt system is further assessed using the Progenetix CNA profiles as input. Firstly, this is done via contrasting the clustering of samples implied by NCIt with an unconstrained clustering. We focus on the 163 level 3 NCIt codes, which are related to 14 “parent” NCIt codes at level 2 of the same hierarchy.

Due to limited data, only the 23 of the 163 level 3 NCIt codes have at least 50 Progenetix samples. For each of these 23 NCIt codes, the average CNA profile was computed. The total dissimilarity (see methods) for these profiles is 83.0. When grouped into 9 clusters based on the NCIt tree the total dissimilarity was reduced to 44.1. Finally, when grouped into 9 clusters based on the unconstrained hierarchical clustering, the resultant dissimilarity was 22.4.

To put these figures in perspective: random partitioning of the 23 profiles into 9 clusters resulted in an average dissimilarity of 55.1. And a dissimilarity of less than 44.1 occurs in about 2% of cases. As such the NCIt partitioning is statistically significant (*p* = 0.02), and the optimal clustering highly significant (*p* < 10^−6^), with a substantially lower total dissimilarity.

The unconstrained clustering is shown in Figure 1. The 9 cluster cutset from the unconstrained clustering has a Rand index (see methods) of 0.77 when compared with the NCIt, indicating moderately strong alignment. The clustering, which is not entirely aligned with NCIt shows how CNA similar samples may occur in diagnostically distinct subsets either because of histopahtological diagnosis does not reflect biology well, or because CNA are not relevant for differentiating mutations.

**Fig 1.**
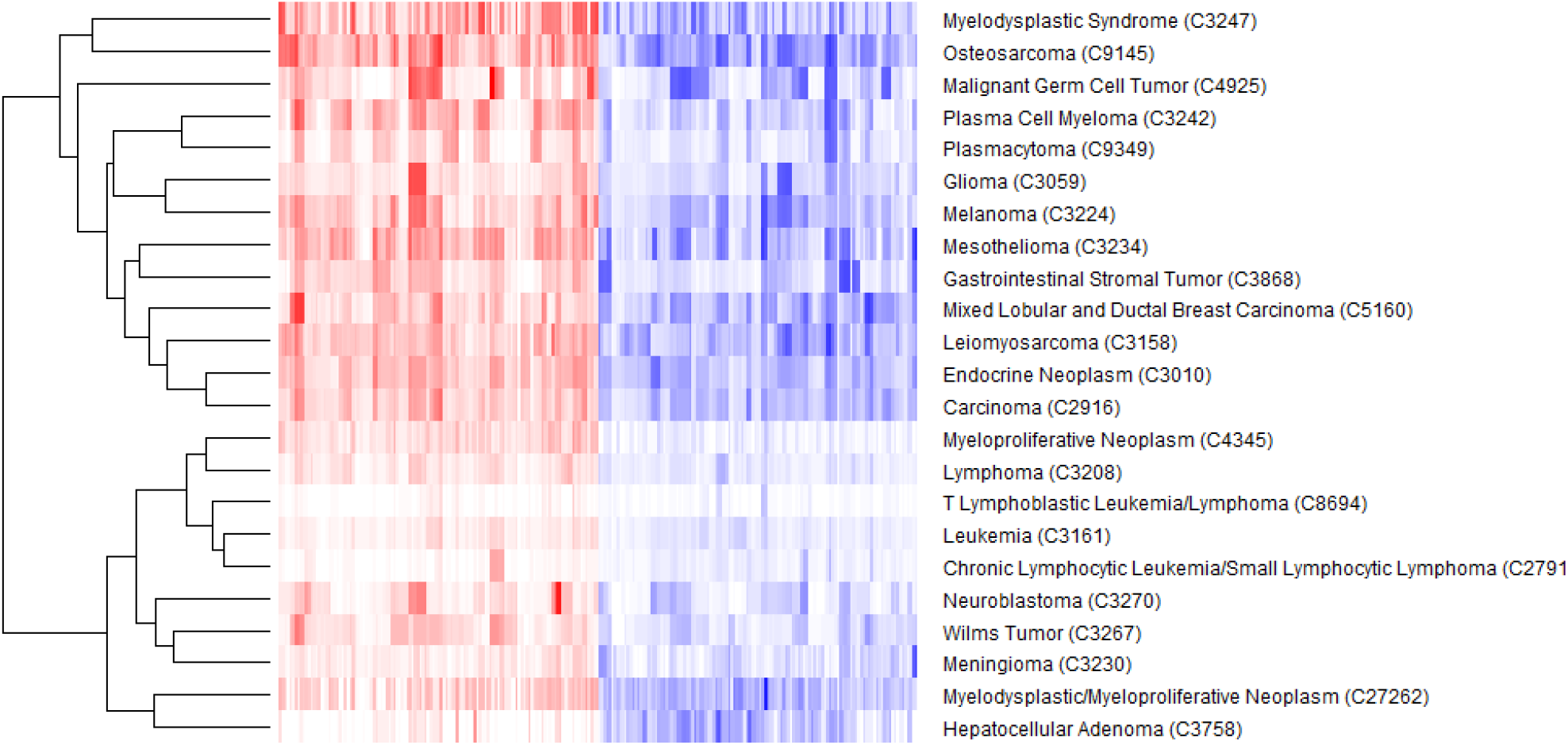
Heatmap of average profiles grouped by unconstrained hierarchical clustering. For each observation vector red and blue cells indicate duplication and deletion CNA respectively.

Qualitatively similar results can be obtained when performing a similar analysis but computing dissimilarity with individual samples rather than averages over groups of samples.

### Sub-clusters in ICD-O groups

Previously we found that the level of detail in the NCIt hierarchy was justified. Here it is tested if additional detail can be justified. This is done by identifying the presence and significance of sub-clusters within ICD-O groups. This is similar to assessing sub-clusters within NCIt groups at a detailed level, but this approach was taken due to a larger number of groups having adequate sample size for analysis.

For the samples for each of the thirty ICD-O cancer types with the highest number of Progenetix samples, the Bernoulli mixture model was fit. Initially the mixture was fit for different number of clusters, and BIC was used to choose the optimal number – where between 4 and 8 clusters were chosen for the different cancers. However, upon visual review, it was not apparent that there were more than 4 distinct clusters in any of the used entity groups. As such, for simplicity, the 4 cluster case was used and presented here.

Regarding interpretation, the greater the reduction in dissimilarity achieved by the clustering, the greater the heterogeneity in the initial single cluster (see methods). There are significantly different sub-clusters apparent in many cases, which cannot be explained by platform.

For each of the 30 ICD-O groups considered the dissimilarity was reduced by more than 11 percent. This was highly significant – e.g., for sample size 1000, and a single cluster with CNA probabilities equal to the CNA probabilities for all Progenetix samples, only 2 percent of 1000 independent replicas obtains a reduction greater 1 percent when fitting 4 clusters. For the three cancers with the most ICD-O samples, their four clusters were visualized according to their CNA probabilities in fig. 3. The four clusters were somewhat similar in shape, or at least sharing some areas with high CNA activity. However the overall level of CNA activity, as well as expression of certain areas, varied across the clusters.

### Clustering of sub-clusters

Given the four estimated sub-clusters per ICD-O, we assessed if the sub-clusters within a cancer were similar, or were rather more alike sub-clusters in other cancer.

For this, for each of the selected 30 ICD-O cancers, for each of its four sub-clusters, the CNA profile was computed – resulting in 120 profiles in total. Next, each profile was compared with all 119 other profiles based on the squared Euclidian distance. The three sub-clusters that were nearest to the selected one were recorded. The hierarchical clustering of the 120 profiles is plotted in fig. 2.

**Fig 2.**
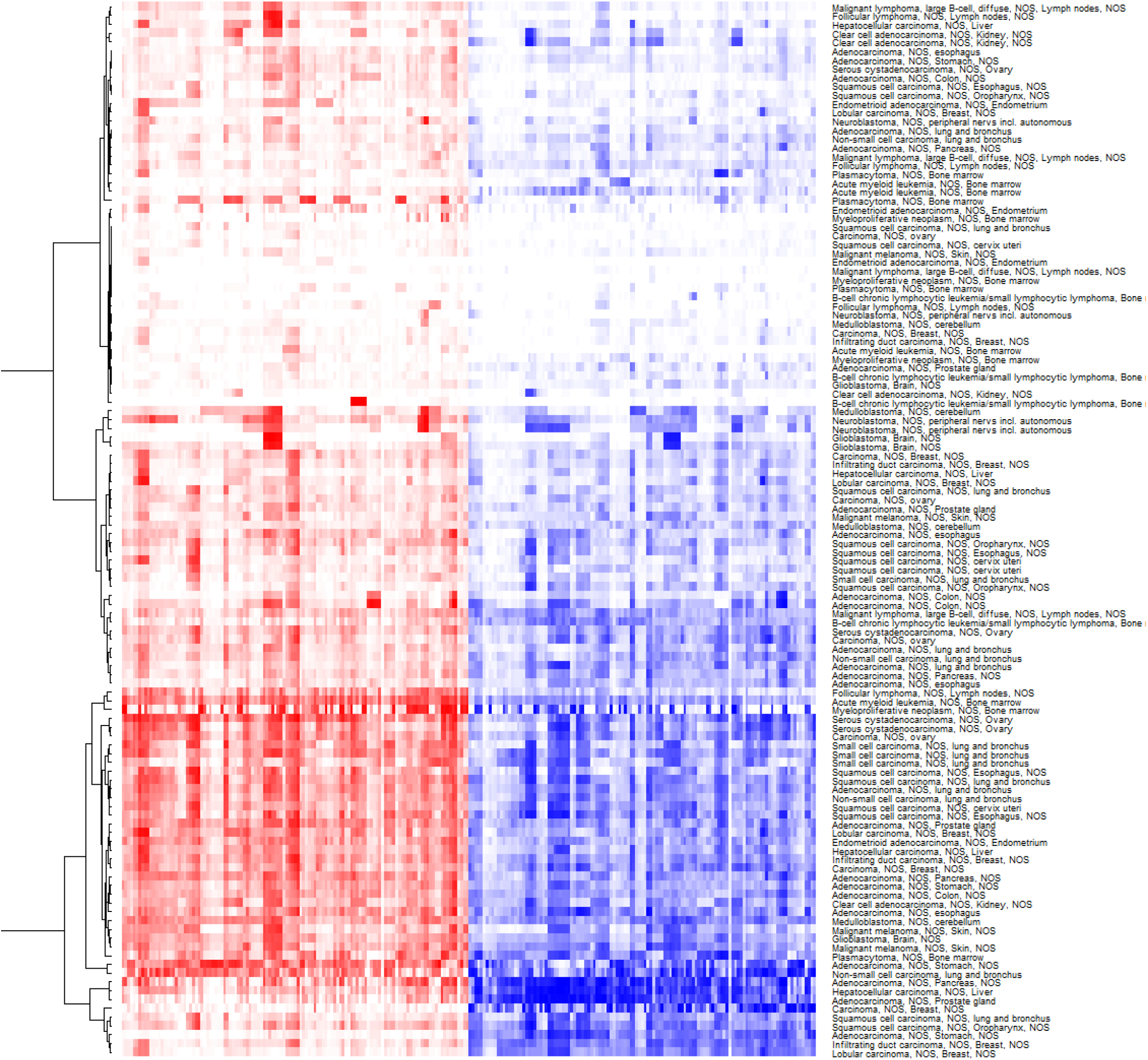
Visualization of the hierarchical clustering of the 120 sub-cluster profiles.

The result of this was that a sub-cluster in a chosen cancer is almost always closer to sub-clusters in other cancers rather than to the other three sub-clusters from that chosen cancer. Specifically, when pooling all 120 sub-clusters together, if one chooses a sub-cluster, on average only 0.18 out of the three sub-clusters that are nearest to it come from the same cancer type. Further, looking at morphology and topography separately, on average only 0.69 and 0.63 of the three nearest pairs had equal ICD-M and ICD-T as the chosen cancer. These observations support the existence of heterogeneity within cancer types and that not all relationships are captured in the current NCIt system or at the current level of detail in ICD-O.

### Exclusion of platform bias

While the Progenetix data is well-suited for large scale analyses of CNV patterns due to its large sample numbers and the limited impact of single-study based biases, in principle analysis results could be skewed due to the inclusion of data from multiple genomic array platforms with possible differences in CNV detection performance and platform derived batches effects should be considered.

For the most represented cancer ICD-O type in Progenetix, 8500/3 C50.9 - Infiltrating ductal carcinoma of the breast, there are 4’633 samples distributed across the 10 array platforms named in Table 4. The Bernoulli mixture model with four clusters was fit to these samples. The profiles of the estimated clusters are shown in fig. 3.

**Fig 3.**
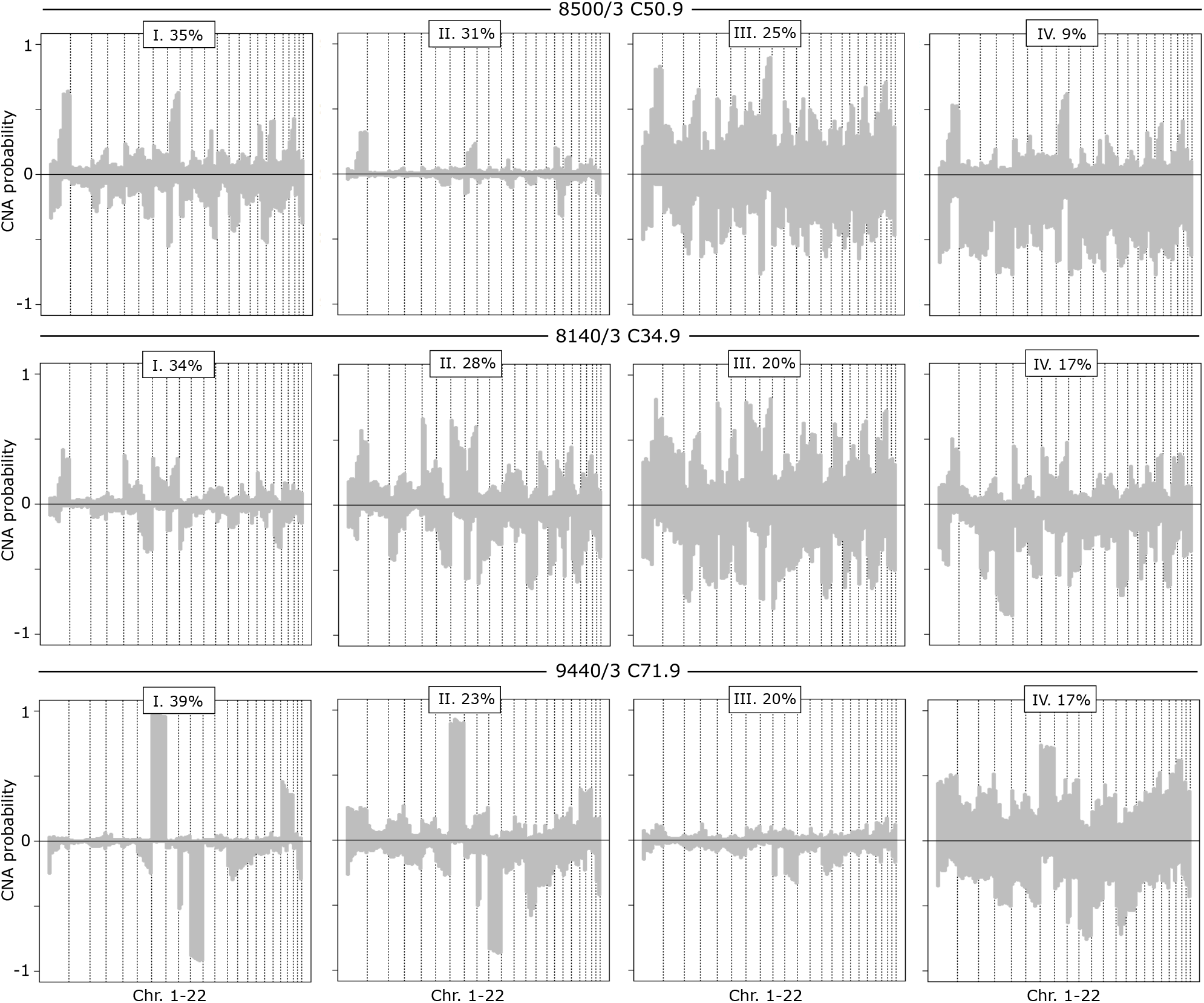
The four clusters for the three ICD-O cancers with the most samples in Progenetix. The clusters are plotted according to their CNA profiles: The average probability of a sample having duplication (positive) or deletion (negative) CNA events along the chromosomes. The percentage of samples in each cluster is given in the box at the top of each sub-plot.

**Table 4.**
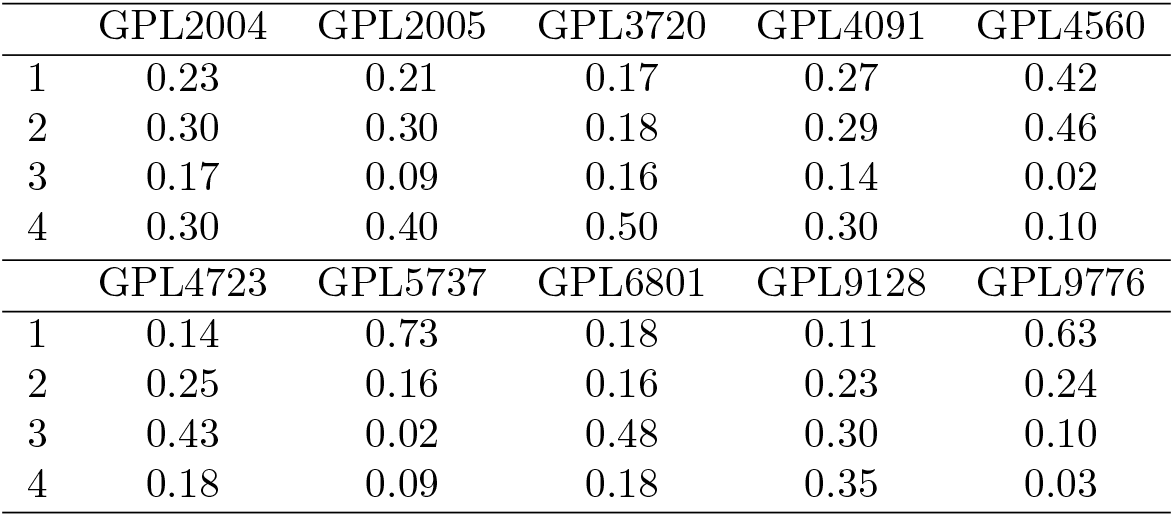
The distribution of 8500/3 - C50.9 (Infiltrating duct carcinoma, NOS - Breast, NOS) cancer samples across the four estimated clusters, conditional on samples coming from a given platform.

|  | GPL2004 | GPL2005 | GPL3720 | GPL4091 | GPL4560 |
| --- | --- | --- | --- | --- | --- |
| 1 | 0.23 | 0.21 | 0.17 | 0.27 | 0.42 |
| 2 | 0.30 | 0.30 | 0.18 | 0.29 | 0.46 |
| 3 | 0.17 | 0.09 | 0.16 | 0.14 | 0.02 |
| 4 | 0.30 | 0.40 | 0.50 | 0.30 | 0.10 |
|  | GPL4723 | GPL5737 | GPL6801 | GPL9128 | GPL9776 |
| 1 | 0.14 | 0.73 | 0.18 | 0.11 | 0.63 |
| 2 | 0.25 | 0.16 | 0.16 | 0.23 | 0.24 |
| 3 | 0.43 | 0.02 | 0.48 | 0.30 | 0.10 |
| 4 | 0.18 | 0.09 | 0.18 | 0.35 | 0.03 |

Visually, the bottom two clusters appear similar, indicating three effective clusters. The distribution of samples across each of the four clusters for a given platform is not exactly uniform (20% in each case), but no clear relation between platform and cluster is apparent. Further assessment of relationship between platform and cluster with other cancers would be challenging due to smaller sample sizes.

## Discussion

CNA profiling data is a powerful measure for meta-analysis since it provides technically consistent whole-genome data. As such, the analysis of extensive amount of CNA profiles can be used as an identifying feature for a given cancer allowing for a generally good representation of cancer entities and the study of its heterogeneity at different levels.

This integrated comparative study is based on a comprehensive subset of copy number data from the Progenetix database. As the largest available collection of this kind of data, it groups studies from different submissions and platforms. If interesting for meta-analysis studies, a series of limitations were also present. First, related to data quality, batch effects can occur as different submissions follow distinct protocols. Furthermore, poor sample annotation and inconsistency, difficulties the treatment and mapping of free diagnosis text into ontology terms. Extensive data curation of samples for harmonization of diagnostic codes and manual mappings between ontologies were required. Other challenges related to the study of the ontological classification arrived from the incomplete (unsampled areas of the hierarchy) and unbalanced (small sample size at the deeper nodes of the hierarchy map) sampling of the hierarchy, which is related to the granularity of samples. To alleviate this, statistical methods were applied to control for unbalanced samples, and mappings to NCIt parent codes allowed for a more consistent level of depth/detail on the hierarchy. As such, to enhance further results, a more strict protocol for biosample submission needs to be adopted, where detailed and standardized characterization of the cancer entities should be required.

The tendency to move from classical classification systems based on pathologic criteria (heavily relaying on the tissue site of origin) to molecular-based taxonomies is still in progress. As cancer stratification is improving, new proposed subclusters are emerging for multiple cancer entities. In this context, not only pathway-based features but “cell-of-origin” characteristics can be useful for classification and understanding of cancers [19]. In the present study, the finding of mixed closeness between the identified subgroups across cancer types could be of interest for further analysis which could guide knowledge transfer from highly studied diseases to poorly understood/rare tumor types.

## Conclusions and future perspectives

In this study, we performed an evaluation of genomic cancer heterogeneity as assessed by the distribution of somatic copy number variation events and the correspondence of emerging subtype clusters with different levels of two common cancer classification systems. Additionally, for the hierarchical NCIt ontology we evaluated the concordance between the classification depth and genomic sample heterogeneity at different classification depth levels.

Whereas current groupings at NCIt are statistically justified, for the detailed NCIt terms as well as for the ICD-O Morphology + Topography pairs some significant heterogeneity and relationships are not captured by the current systems. Further statistically significant sub-clusters were found for individual terms, highlighting the need of improved classification systems. A further stratification of cancer entities could hopefully be achieved through the wider adoption of molecular classifiers such as has been shown previously in medulloblastomas [6] or renal cell carcinomas [20], among others. For data integration purpose and fidelity of the results, we stress the need of improved systems for standardized submission of sample information, especially considering ontology based classifications and data formats supported through initiative such as GA4GH and HL7 [21, 22].

