## Supplemental Table 1 & 2 for "Copy number variant heterogeneity among cancer types reflects inconsistent concordance with diagnostic classifications"

### Supplementary material

**Table 1.** Average coverage, and standard deviation, for ICD-O pairs.

| Cancer type (ICD-O M-T) | Mean | Standard deviation |
|---|---|---|
| 8010/3 - C56.9 | 0.28 | 0.23 |
| 8046/3 - C34.9 | 0.29 | 0.21 |
| 8070/3 - C10.9 | 0.27 | 0.2 |
| 8070/3 - C34.9 | 0.37 | 0.23 |
| 8120/3 - C67.0 | 0.23 | 0.17 |
| 8140/3 - C16.9 | 0.2 | 0.22 |
| 8140/3 - C18.9 | 0.25 | 0.19 |
| 8140/3 - C34.9 | 0.29 | 0.2 |
| 8140/3 - C61.9 | 0.18 | 0.21 |
| 8170/3 - C22.0 | 0.29 | 0.23 |
| 8310/3 - C64.9 | 0.21 | 0.19 |
| 8380/3 - C54.1 | 0.16 | 0.22 |
| 8441/3 - C56.9 | 0.53 | 0.23 |
| 8500/3 - C50.9 | 0.25 | 0.21 |
| 8720/3 - C44.9 | 0.29 | 0.19 |
| 9440/3 - C71.9 | 0.24 | 0.18 |
| 9470/3 - C71.6 | 0.21 | 0.2 |
| 9500/3 - C47.9 | 0.16 | 0.18 |
| 9680/3 - C77.9 | 0.12 | 0.13 |
| 9690/3 - C77.9 | 0.08 | 0.11 |
| 9731/3 - C42.4 | 0.17 | 0.19 |
| 9823/3 - C42.4 | 0.03 | 0.08 |
| 9835/3 - C42.4 | 0.09 | 0.13 |
| 9861/3 - C42.4 | 0.03 | 0.07 |
| 9989/3 - C42.4 | 0.11 | 0.16 |

**Table 2.** NCIt Morphology tree partitioned into its 14 level 2 cutsets. The first column is the name of the level 2 NCIt node whose cutset (progeny) is taken. The second column is the number of unique NCIt codes in the cutset, and third the number of Progenetix samples corresponding to those NCIt codes.

| NCIt | N. of NCIt child | N. of samples |
|---|---|---|
| C3709 | 661 | 33114 |
| C3708 | 25 | 289 |
| C7069 | 10 | 0 |
| C27134 | 312 | 7931 |
| C7058 | 21 | 1637 |
| C6971 | 19 | 280 |
| C7059 | 100 | 656 |
| C3786 | 18 | 78 |
| C6930 | 63 | 306 |
| C6974 | 5 | 0 |
| C7068 | 7 | 0 |
| C35562 | 115 | 5316 |
| C7061 | 13 | 0 |
| C3422 | 7 | 0 |
